# Scaling of the Bicoid morphogen gradient: the effect of state dependent diffusion

**DOI:** 10.64898/2025.12.15.694169

**Authors:** Priya Chakraborty, Shyam Iyer, Richa Rikhy, Mithun K. Mitra, Amitabha Nandi

## Abstract

The mechanisms underlying the scaling of the Bicoid morphogen gradient in *Drosophila* with the embryo size is not clearly understood. We propose a model with a spatially varying diffusion coefficient along the anterior-posterior axis, that is proportional to the nearly periodic spatial distribution of the nucleo-cytoplasmic domains and alternating between regions of fast and slow diffusion. We postulate that for specific interpretation of the heterogeneous environment, where the space available for free diffusion within an energid is assumed to be proportional to the embryo size, a change in the embryo size can lead to a size-dependent scaling of the gradient lengthscale. We further study a two component model with slow and fast diffusing Bicoid, and identify this model to be equivalent to the heterogeneous diffusion model further postulating that the *fast-state occupancy* which is a measure of the fraction of time spent in the fast diffusing state, should also scale with the embryo size via the energid size. Finally, we incorporate nuclear shuttling into our model to understand the effect of shuttling on the gradient lengthscale and scaling with embryo size. We argue that for the particular case where the degradation within the nucleus is low, nuclear shuttling does not perturb the Bicoid gradient. Our study suggests that scaling of the gradient length scale is possible due to spatial heterogeneity and does not depend on nuclear trapping as suggested previously.

## I. INTRODUCTION

The non-uniform distribution of the morphogen in a developing tissue provides crucial positioning information and controls subsequent decisions about cell fate [1]. Bicoid, a homeodomain transcription factor, is one of the most extensively studied and first experimentally verified morphogens that provide positional information to multiple zygotic genes (GAP, Hb, etc.) in the early stages of *Drosophila* embryogenesis [2–4]. In the simplest theoretical model, known as the Synthesis, Diffusion, and Degradation model (SDD), the Bicoid protein that gets *synthesized* from the maternally deposited Bicoid mRNA in the anterior pole of the *Drosophila* embryo (with a flux *Q*), *diffuses* from the source (with diffusivity *D*) along the anterior-posterior (AP) axis, accompanied by a uniform *degradation* rate (degradation coefficient *µ*) [5, 6], to establish a concentration gradient of the Bicoid protein. Due to the interplay between production at the source, diffusion, and the degradation rate, a steady state is achieved and an exponentially decaying concentration gradient profile 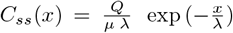, is established along the AP axis with a characteristic lengthscale 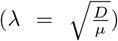 Subsequent readout of the Bicoid concentration by the downstream genes defines the developmental pathway.

Several models have been proposed in recent years to address the limitations of the SDD model. The nuclear shuttling model [7], the non-localized mRNA source model [8], and the multiple dynamic modes model (fast-slow model) [9] represent notable approaches. In the dynamic cytoplasmic environment of a syncytial Drosophila embryo, the number of nuclei multiplies in each nuclear division cycle (cycle 1 − 14) during the establishment of the Bicoid gradient. In the nuclear shuttling model, the concentration profile is established through the Bicoid diffusion along with nucleocytoplasmic shuttling in the presence of the growing number of nuclei [7]. In [8], Cheung et al. proposed that the adaptability of Bicoid gradient profiles in Drosophila embryos is achieved through spatial and temporal regulation of mRNA sources, ensuring robust embryonic patterning despite environmental fluctuations. More recently, Athilingam et al. described a model [9] incorporating multiple dynamic modes of Bicoid, distinguishing between fast and slow diffusion processes to explain the long-range Bicoid pattern and intricate dynamics involved in Bicoid gradient formation.

An important feature of the Bicoid gradient is its scalability. Embryos can vary significantly in size both within and between species, but, interestingly the organism seems to sense this size variability and adjust the characteristic length-scale accordingly. This ability to scale is crucial, as it allows target genes to read positional information and appropriately set the gene expression boundaries in the right relative positions, ensuring proper body patterning and axis formation during the next stage of development. In the case of Drosophila, the segmented structure of the adult organism maintains consistent proportions, even across an extensive range of embryo sizes (400–600*µm*) [10]. However, despite its simplicity and its ability to model the formation of the Bicoid gradient in the syncytial embryo, the SDD model does not account for the invariance in length scaling across embryos of different sizes. Additionally, experimental measurements of the effective diffusion coefficient of Bicoid vary widely, from 0.2 to 18 *µm*^2^*/s* [7, 11]. A previous study by Gregor *et al*. [6] had suggested a plausible mechanism for scaling of the characteristic lengthscale, by incorporating nuclear shuttling, showing that the characteristic length scales linearly with the system size *L* (i.e. *λ* ∼ *L*). However, the model relies heavily on the nuclear trapping dynamics with the important assumption that the entire degradation is considered to take place mainly within the nucleus. This mechanism is thus at odds with the important experimental observation that the characteristic lengthscale is independent of nuclear trapping and high diffusivity of Bicoid can establish a stable gradient by diffusion before cycle 8 [12, 13].

The SDD model, or the subsequent variants, do not explicitly address the impact of the dynamic, heterogeneous environment on Bicoid diffusion, particularly in the context of scaling of gradient length scale with embryo size. The internal micro-environment of a developing syncytial embryo is highly heterogeneous. Starting from oviposition, the nuclei proliferate up to cell cycle 8 in the central cytoplasmic region, and at cell cycle 9, the nuclei migrate to the embryo’s surface region. Here, the cytoplasmic island divides into nuclear/cytoplasmic protoplasmic islands, also known as energids. Previously, it was argued that in the syncytial Drosophila embryo with periodically occurring energids, the morphogen may exhibit rapid diffusion within an individual energid but only slow diffusion between energids, as demonstrated by photobleaching experiments [14]. In some recent studies, gradients of photo-activated [15] and plasma-membrane binding proteins [16] were quantified and showed that the spatial heterogeneity induced by the presence of the dynamic cytoarchitecture together with the periodic array of energids play an important role in shaping a steady state morphogen gradient. In a more recent study [9], the authors suggested that Bicoid can have multiple dynamic modes of fast and slowly diffusing species in order to achieve the observed AP patterning. However, no model has explored the effect of periodically occurring energid structure on modifying diffusion coefficient and further length scaling in a Drosophila embryo.

Motivated by these recent developments, in this paper, we theoretically study the effect of heterogeneity on the scaling properties of the Bicoid gradient. We propose a model where the heterogeneous environment inside the embryo is manifested in a diffusion coefficient periodically varying in space (see Fig. 1(a)). We show that the spatial heterogeneity in diffusion can lead to specific conditions for scaling of the characteristic lengthscale with change in embryo length. We further compare this heterogeneous diffusion model with the previously studied multiple dynamic mode model [9] (see Fig. 1(b)), and assuming detailed balance in the kinetics of switching between the two modes, we draw an equivalence between the two models. We further study the effect of nuclear shuttling in our proposed model and discuss the relevance of the present work on the scaling of the Bicoid gradient and future perspectives. In this study, we use experimentally estimated parameters, which are listed in Table I in Appendix A.

**TABLE I.** Parameter values for the different models.

| Parameter | Physical meaning | Value | Ref. |
| --- | --- | --- | --- |
| $D_1(D_f)$ | Fast diffusion coefficient in the heterogeneous diffusion model (Diffusion coefficient of fast bicoid species in multiple dynamic mode) | $18 \mu m^2.s^{-1}$ | [11] |
| $D_2(D_s)$ | Slow Diffusion coefficient in the heterogeneous diffusion model (Diffusion coefficient of slow bicoid species in multiple dynamic mode) | $0.3 \mu m^2.s^{-1}$ | [6] |
| $l_1$ | Length of region for fast diffusion in the heterogeneous diffusion model | $7 - 12 \mu m$ | [16] |
| $l_2$ | Length of region for slow diffusion in the heterogeneous diffusion model | $1 \mu m$ | [16] |
| $\beta_s$ | Transition rate of slow to fast bicoid species | $0.02 s^{-1}$ | [9](Model 3 in Table. S7) |
| $\beta_f$ | Transition rate of fast to slow bicoid species | $0.07 s^{-1}$ | [9](Model 3 in Table. S7) |
| $\mu$ | Degradation coefficient | $0.00035 s^{-1}$ | [9] |
| $\beta_c^f$ | Rate of unbinding of the fast bicoid species from nucleus | $0.014 s^{-1}$ | [6, 29] |
| $\beta_N$ | Rate of forward nuclear trapping | $0.0028 s^{-1}$ | [7] |
| $\beta_c^s$ | Rate of unbinding of the slow bicoid species from nucleus | -- | -- |

**FIG. 1.**
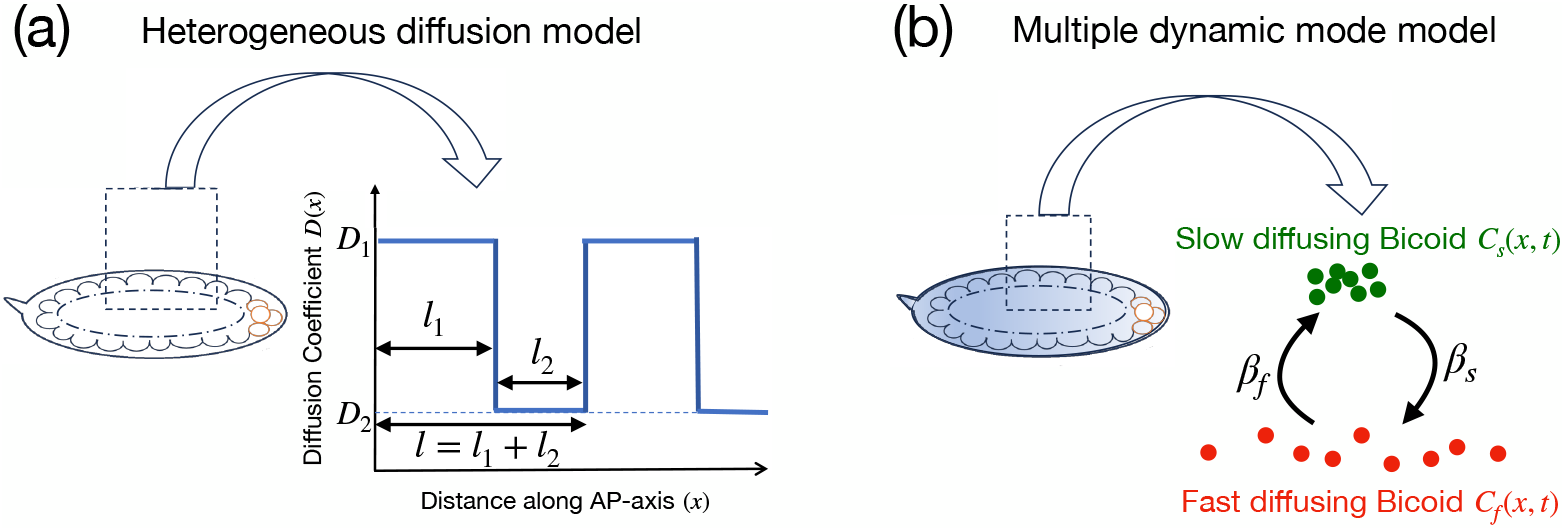
Schematic representation of the different models of Bicoid diffusion studied in this paper. (a) Heterogeneous diffusion model with the diffusion coefficient proportional to the periodicity of the nucleo-cytoplasmic domains along the AP-axis and alternating between regions of fast and slow diffusion. An equivalent representation is: (b) a multi-state dynamic model in which Bicoid consists of two interconverting subpopulations: a slow-diffusing species (green filled circles) and a fast-diffusing species (red filled circles).

## II. HETEROGENEOUS DOMAIN MODEL

Bicoid diffuses along the cortical region of the Drosophila embryo to establish a gradient. While most existing models in the literature assume a homogeneous environment in this cortical region, the underlying cytoplasmic, cytoskeletal and nuclear distribution suggests that the homogeneity assumption may need to be re-examined. The energids which compartmentalize the nuclei are partially separated by the membrane furrows, which extend vertically into the nuclear interior during interphase. Thus, for cytoplasmic Bicoid diffusing in this furrow regions, the diffusion is likely to be constrained as compared to the non-furrow regions. Similar heterogeneities can also occur from other biophysical causes. For example, the nuclear volume also constrains diffusion compared to non-nuclear spaces, and this can once again lead to spatially heterogeneous diffusion. During early cellularisation, the tubulin cytoskeleton provides a structural scaffold that further compartmentalises the cytoplasm and creates segregated nuclear units. In addition, the flows originating from the central region of the embryo’s cortex during early development can also give rise to transient, energid-like domains, further contributing to local heterogeneity in the diffusion landscape. The effect of such periodic spatial heterogeneity and its consequent effect on gradient formation has not been investigated.

We propose a periodic two-domain model for Bicoid diffusion, with the periodicity set by the energid lengthscales. The time evolution of the Bicoid concentration is then described by

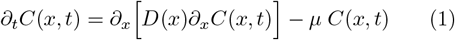

with the periodic diffusion coeficient given by

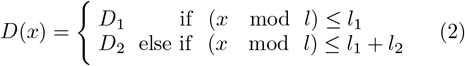

where the total length of the energid is *l* = *l*_1_ + *l*_2_. The region of length *l*_1_ corresponds to the fast diffusion, while *l*_2_ corresponds to slow diffusion, *D*_1_ *> D*_2_ (Fig. 1(a)). This spatially varying diffusion coefficient is then periodically repeated. The total length of an embryo thus can be represented as *L* = *n l*, where *n* is the number of energids along the AP axis. Fig. 1(a) shows a schematic diagram of the model consideration. We assume that the degradation is uniform across the length of the embryo, and Bicoid production happens through a point source at the anterior pole (*x* = 0).

This one-dimensional model with a periodically varying diffusion coefficient has been proposed and investigated in the literature, dating back to the Maxwell Garnett theory of inclusions in an optical medium [17]. An effective medium theory can be written where Eq. 1 can be replaced by the following time-evolution equation for the Bicoid concentration *C*(*x, t*)

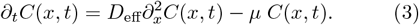

Here *D*_eff_ is the effective diffusion coefficient: an exact analytic expression for *D*_eff_ can be obtained either from the forward formalism [18], by directly using Eq. 1, or using the backward formalism [19], both of which gives,

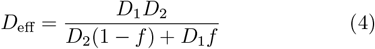

where, *f* = *l*_2_*/*(*l*_1_ + *l*_2_) is the fraction of the energid length occupied by the slow diffusing region.

We compare the steady concentration profile *C*_*ss*_(*x*) obtained by numerically solving Eq. 1 at steady state (see Appendix B), to the steady-state concentration profile obtained by solving Eq. 3, which is given by 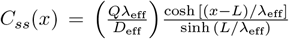. Here 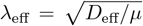 is the corresponding characteristic length-scale (see Appendix B). We compare the two steady-state profiles in Fig. 2 for *n* = 50. As can be seen from the figure, with increasing number of energids (see Inset of Fig. 2 for *n* = 1, 2, and 10), the effective diffusivity model is in very good agreement with the full heterogeneous diffusion model.

**FIG. 2.**
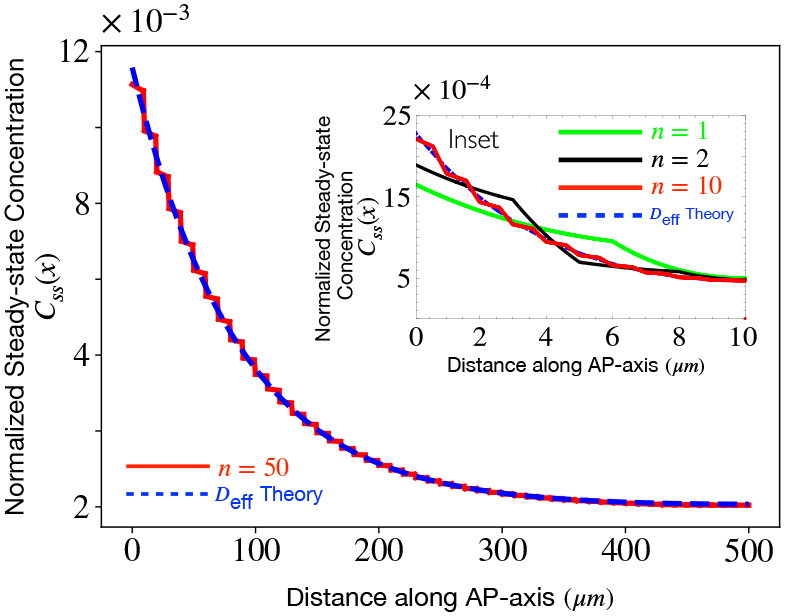
Normalized steady state Bicoid concentration profile due to heterogeneous diffusion. The solid line (red) represents the gradient obtained by numerically solving Eq. 1 at steady state, and the dashed line (blue) represents the corresponding steady state gradient with an effective diffusion coefficient (Eq. 3). Parameter values used are *D*_1_ = 18 *µm*^2^.*s*^−1^, *D*_2_ = 0.3 *µm*^2^.*s*^−1^, *µ* = 3.5 *×* 10^−4^ *s*^−1^, *l*_1_ = 9 *µm, l*_2_ = 1 *µm, L* = 50 *µm* for an energid number *n* = 50. The Inset shows a comparison between the original model (Eq. 1) and the effective model (Eq. 3) as a function of energid number. The parameters for the Inset are in arbitrary units and does not correspond to the biologically relevant parameters; they are chosen to be *D*_1_ = 1, *D*_2_ = 0.2, *µ* = 0.02, *L* = 10. With an increase in energid number *n*, the agreement becomes better.

We now investigate whether this heterogeneous domain model can yield a scaling of the gradient length scale with embryo size. In this formalism, an increase in the total length of the embryo can be modeled through three different microscopic scenarios – (i) With increasing length, the number of energids *n* increases, with *l*_1_ and *l*_2_ remaining unchanged, (ii) The number of energids remains unchanged, but the size of the energid increases proportionately such that the ratio *l*_1_*/l*_2_ remains the same, and (iii) again, as in (ii), the number of energids remains unchanged and the total energid size increases proportionately, but this increase is only due to the increase of either of *l*_1_ or *l*_2_, such that the ratio *l*_1_*/l*_2_ changes.

Recent experiments focusing on the nuclear degradation of Bicoid suggests that the number of nuclei does not change with the change in sizes of embryos [6, 11]. This implies that scenario (i) as proposed above is not biologically relevant, and thus only the second and third possibilities can be true with increasing length of the embryo. Since the effective diffusion constant equation depends on the ratio *l*_1_*/l*_2_ when *D*_1_, *D*_2_ remain unchanged, for the gradient length to scale with embryo size, the only possibility is that the individual domain lengths *l*_1_ and *l*_2_ change with embryo length in the energid. We choose the fast and slow diffusion coefficients to the extremes of the observed Bicoid diffusivities, *D*_1_ = 18 *µm*^2^.*s*^−1^, *D*_2_ = 0.3 *µm*^2^.*s*^−1^. *D. melanogaster* embryos have typical lengths around 500*µm* with a spread ranging from 400 − 600*µm* [10, 20, 21]. We choose the number of energids *n* = 50 and vary *l*_1_*/l*_2_ in the range 7−12 to capture this variation in embryo lengths [16]. The effective diffusion coefficient for these parameters is obtained from Eqs. (4) and is shown in Fig. 3(a) (red line). The corresponding scaling of the effective characteristic length-scale (*λ*_eff_) with the change in embryo length is shown in Fig. 3(a) (blue line). As can be seen from the figure, the heterogeneous domain model corresponding to the scenarios where the lengths of the two domains *l*_1_ and *l*_2_ change individually with increasing embryo length is sufficient to reproduce a scaling of the Bicoid gradient. This ensures that the concentration across the segmented body structure is proportionally maintained, allowing the downstream genes to interpret the positional information accurately. Moreover, since the effective diffusion coefficient *D*_eff_ is independent of *n*, the scaling behavior predicted by this heterogeneous model remains consistent across nuclear cycles, further supporting its biological relevance.

**FIG. 3.**
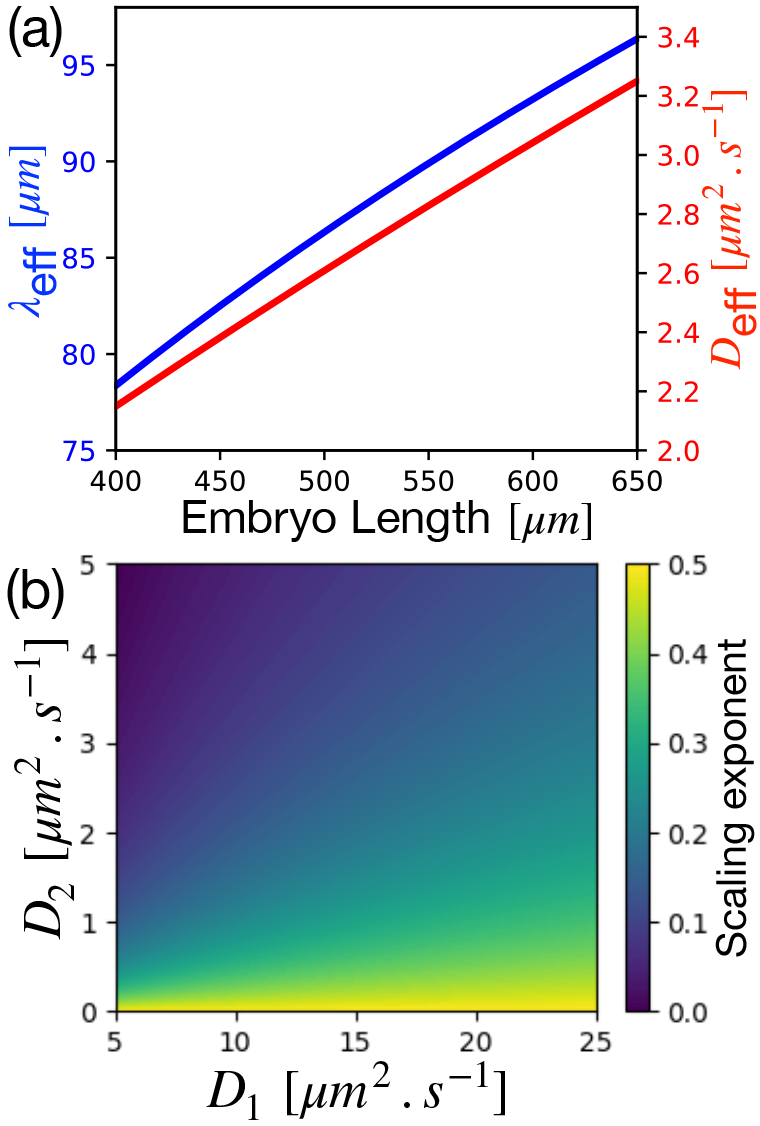
Scaling of the characteristic lengthscale *λ*_eff_ for the heterogeneous diffusion model with embryo length *L* (a) *D*_eff_ (red) and *λ*_eff_ (blue) plotted versus embryo length *L* for *D*_1_ = 18 *µm*^2^.*s*^−1^, and *D*_2_ = 0.3 *µm*^2^.*s*^−1^. The scaling exponent *α* in this case is *α* ≈ 0.425. (b) Colormap showing the scaling exponent *α* in the (*D*_1_–*D*_2_) plane. The range of the exponent values obtained varies in the range [0−0.5]. For both the plots, the values of the other parameters are *µ* = 3.5*×*10^−4^ *s*^−1^, *l*_2_ = 1 *µm*, and *n* = 50. Here *l*_1_ is varied from 7 *µm* to 12 *µm*. For (b)*µ* = 3.5 *×* 10^−4^.

Using the effective representation of the heterogeneous diffusion model discussed above, we now proceed to study the scaling of the characteristic lengthscale *λ*_eff_ with embryo length *L*: we assume *λ*_eff_ ∝ *L*^*α*^ and try to estimate the exponent *α* value at the different limits. From Eq. 4 we note that when *D*_1_*/D*_2_ ≫ 1, we obtain, *λ*_eff_ ∼ *L*^1*/*2^, while in the limiting case, *D*_1_*/D*_2_ = 1, we recover, *λ*_eff_ ∼ *L*^0^, the classical SDD result where there is no scaling of the length. Thus the scaling exponent *α* varies in the range [0 − 0.5] depending on the ratio of the fast and slow diffusion. In Fig. 3(a) we show the scaling of *λ*_eff_ (and *D*_eff_) with the embryo length *L* for a scenario where *D*_1_*/D*_2_ is large. Furthermore to show the range of scaling exponents *α* obtained by varying the *D*_1_*/D*_2_ ratio, we fit *λ*_eff_ ∝ *L*^*α*^, and the exponent *α* for different combinations of the diffusion coefficient *D*_1_ and *D*_2_ is shown in a phase plot in Fig. 3(b).

## III. DIFFUSION MODEL OF BICOID WITH MULTIPLE DYNAMIC MODES

Previous experiments has reported a diffusion coefficient of Bicoid ≤ 1 *µm*^2^*/s* [6]. The timescales of pattern formation in the early Drosophila embryo (within ∼ 1 hour of fertilization) is at odds with this low value of the Bicoid diffusivity. Recent developments in the field suggest that there can be more than a single dynamic mode for the diffusion of Bicoid protein to establish anterior-posterior patterning [9]. Further, in photobleaching studies, it is observed that all recovery curves begin at a level significantly higher than the background. Thus, a portion of the Bicoid may already be re-equilibrated, indicating the possibility that a part of the Bicoid has a higher diffusion coefficient than the rest [4]. Together, these studies suggest that the Bicoid protein may be in one of two states - a fast state or a slow state, with a corresponding diffusion constant, along with a rate of interconversion between the two states. Fig. 1(b) shows a schematic representation of the model consideration.

We consider a two species model, with the time-evolution of the slow and fast Bicoid species *C*_*s*_(*x, t*) and *C*_*f*_ (*x, t*) respectively given by

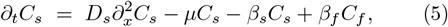

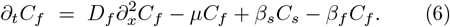

Here, *D*_*s*_ and *D*_*f*_ are the diffusivities of the slow and fast Bicoid species respectively. The transition rates (in dimensions of inverse time) from slow to fast species and vice-versa are denoted as *β*_*s*_ and *β*_*f*_ respectively. Equivalently, one can describe the kinetics by the *mean lifetime* of the slow (fast) diffusing state 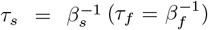: henceforth, we will use both the definitions interchangeably for convenience. The degradation rate *µ* is assumed to be same for both for the slow and fast. The diffusivities and the transition rates are assumed to be independent of position, and hence this can be considered as a simplified version of the model discussed in [9]. Within this two species model, we can determine the effective diffusivity, and hence the characteristic lengthscale of the Bicoid gradient.

We make an adiabatic elimination by assuming a detailed balance condition *β*_*s*_*C*_*s*_ = *β*_*f*_ *C*_*f*_ . This assumption is reasonable if the average residence times in each state (i.e. slow or fast) are much smaller than the characteristic diffusional transport timescales i.e. 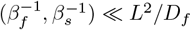. We note that the diffusional timescale *L*^2^*/D*_*f*_ ∼ 10^4^ *s* (assuming *L* ≈ 500*µm* and *D*_*f*_ ≈ 20*µm*^2^.*s*^−1^). We further note from Ref. [9] (see Table. S7), that the mean residence times ∼10^2^ *s* are significantly smaller that the diffusional timescales, justifying our approximation.

We define the *fast-state occupancy* as 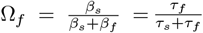, a dimensionless parameter which is a measure of the steady-state relative occupancy in the fast-diffusing state (0 *<* Ω_*f*_ *<* 1) and is more informative than the individual rates. We therefore obtain the time-evolution equation for the total Bicoid concentration *C*_*T*_ (*x, t*) = *C*_*s*_(*x, t*) + *C*_*f*_ (*x, t*)

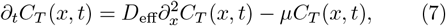

where *C*_*s*_(*x, t*) = (1 − Ω_*f*_ )*C*_*T*_ (*x, t*), and *C*_*f*_ (*x, t*) = Ω_*f*_ *C*_*T*_ (*x, t*). Eq. 7 leads to the same steady-state solution as Eq. 3 with the characteristic length-scale 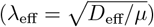 now defined in terms of new effective diffusion coefficient *D*_eff_ which is given as

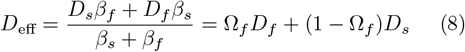

From Eq. 8, we note that the effective diffusivity is a function of the fast-state occupancy Ω_*f*_ .

### A. Equivalence between models

We see that both the heterogeneous diffusion model and the diffusion model with multiple dynamic modes predicts an effective diffusion coefficient for bicoid from the underlying microscopics. Comparing the *D*_eff_ given by Eq. 4 and Eq. 8, we can draw an equivalence between our proposed heterogeneous diffusion model and the two-state model of Bicoid diffusion.

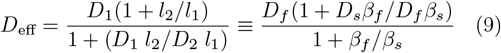

Identifying *D*_1_ = *D*_*f*_ and *D*_2_ = *D*_*s*_, the above equivalence therefore will hold true if the ratio of the switching rates satisfy

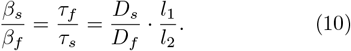

For the heterogeneous diffusion model the typical time spent in the compartment of size *l*_*i*_ or the typical time taken to exit the box is given by 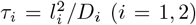. Eq. 10 can then be rewritten as 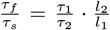. We note that for the equivalence between the models to hold, the switching rates in the two-state Bicoid model are not simply equal to the inverse typical times in the heterogeneous diffusion model. An absolute magnitude of the rates can be defined by introducing a weight factor proportional to the relative size of each compartment, i.e.

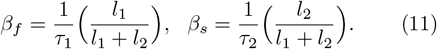

With this identification for the absolute rates, we obtain, *β*_*f*_ ≈ 0.18 *s*^−1^ and *β*_*s*_ ≈ 0.02 *s*^−1^ (taking *l*_1_ = 9*µm* and *l*_2_ = 1*µm*) which are similar to the values reported in [9] (also see Table I in Appendix A) within the experimental uncertainties. Within this framework, the two-species model can be interpreted as a systematic coarse-grained continuum representation of the underlying microscopic structure which leads to slow (fast) diffusion. Eq. 10 can be further rewritten to obtain an expression for the fast-state occupancy

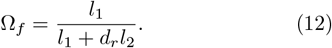

Here 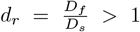 is the ratio of the fast and slow diffusivities and is held fixed in our study. Note that the fast-state occupancy Ω_*f*_ is the natural control parameter for the two-state Bicoid model and by looking at Eq. 12, we thus note that an increase in the length *L* of the embryo by the heterogeneous diffusion model (Sec. II), leading to an increase in the free energid space (by increasing *l*_1_) can equivalently lead to an increase in the fast-state occupancy Ω_*f*_ . This can be interpreted as a decrease (increase) in the conversion rate of fast- to slow-state Bicoid (or conversely slow- to fast-state) species. The regulation of the effective diffusion coefficient (*D*_eff_) and the characteristic lengthscale (*λ*_eff_) with an increase in Ω_*f*_ is shown in Fig. 4. At large lengthscales then, both of the heterogeneous diffusion and the two species model can explain the scaling of the gradient, independent of the the origin of the heterogeneity - whether structural (two-domain), or chemical (two-species). In Fig. 4, we show the scaling of the characteristic lengthscale; instead of directly plotting it versus the embryo length, we plot is as a function of Ω_*f*_ which scales as ∼ *l*_1_(*L*) (see Eq. 12).

**FIG. 4.**
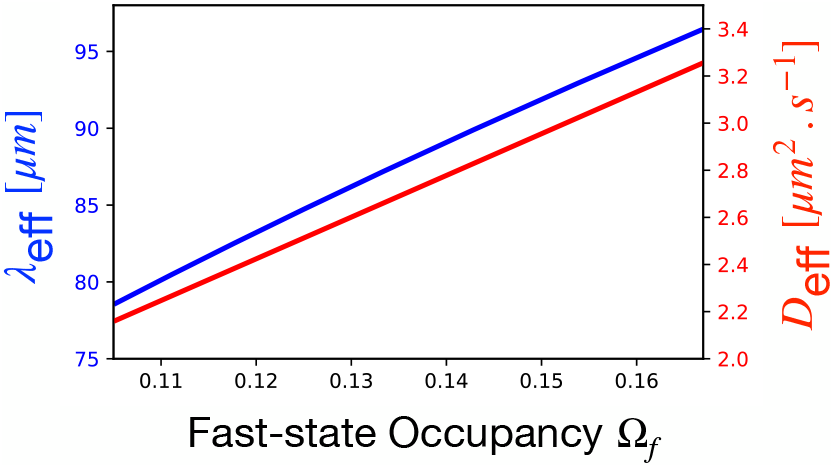
Scaling behavior of the characteristic lengthscale (blue) and the effective diffusion coefficient for the two Bicoid species model discussed in Sec. III. To show the equivalence of this model with the heterogeneous diffusion model, instead of the embryo length *L*, here we vary Ω_*f*_ which is a function of *L* via Eq. 12. The fixed parameter values are *µ* = 3.5 *×* 10^−4^ *s*^−1^, *D*_*f*_ = 18 *µm*^2^.*s*^−1^, and *D*_*s*_ = 0.3 *µm*^2^.*s*^−1^. The scaling exponent in this case is *α* ≈ 0.445

Here, we have argued that this simple fast-slow model, under an internal chemical equilibrium assumption, is equivalent to the more complicated heterogeneous diffusion model and can account for the scaling of characteristic lengthscale *λ*_eff_ with embryo length *L* provided the fast-state occupancy Ω_*f*_ satisfies Eq. 12. In Appendix C, we also derive the leading order corrections to the gradient lengthscale if the diffusivities of the slow and fast species have a linear dependence on the distance from the anterior pole [9], and show that this provides only weak corrections to the results derived here. The consideration of an increase in the lifetime of the Bicoid species in the fast state relative to the slow state equivalently scales the effective diffusion coefficient and characteristic lengthscale similar to the heterogeneous diffusion model. The consideration of an increase in energid length by an increase in matrix length *l*_1_, keeping the inclusion length *l*_2_ fixed, specially allocates a greater region of higher diffusivity, that is physically equivalent to an increase in the Ω_*f*_ value of the Bicoid species. This shows that multiple dynamic modes of Bicoid allow not only long range pattern formation but also enable length invariability within and across species.

## IV. MULTIPLE DYNAMIC MODES WITH NUCLEOPLASMIC SHUTTLING

Having argued that the two species Bicoid model is a simpler representation of the heterogeneous diffusion model, we now use it to study the effect of nuclear trapping of Bicoid on the scaling properties. Apart from the slow and fast Bicoid species *C*_*s*_ and *C*_*f*_, we now consider an additional species *C*_*b*_ representing the nuclear-bound Bicoid. Thus, the total Bicoid concentration is now given by *C*_*T*_ = *C*_*free*_ +*C*_*N*_ = *C*_*f*_ +*C*_*s*_ +*C*_*b*_, where *C*_*free*_ = *C*_*f*_ +*C*_*s*_ and the time evolution Eqs. (5–6) with the inclusion of the bound species *C*_*b*_ modifies to

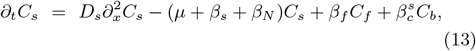

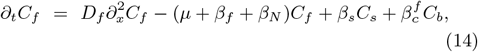

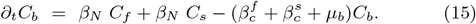

Here *µ*_*b*_ is considered as the degradation rate for the nuclear-trapped Bicoid which is considered to be different from degradation rate *µ* for the free species. Here 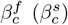 is the rate of unbinding from the nucleus for the fast (slow) Bicoid and *β*_*N*_ is the rate of forward nuclear trapping.

As before we assume local detail balance for the chemical kinetics between the fast and the slow species of the Bicoid, i.e. *β*_*s*_*C*_*s*_ = *β*_*f*_ *C*_*f*_ . This leads to further simplification allowing us to express the bound Bicoid concentration *C*_*b*_ in terms of the total free species concentration *C*_*free*_.

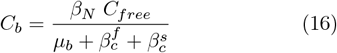

Using Eq. 16, the differential equation for 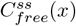 at steady state is given by Eqs. 17

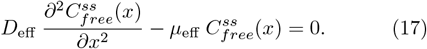

Here the effective diffusion coefficient *D*_eff_ and effective degradation constant *µ*_eff_ are as follows.

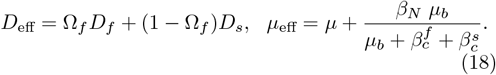

We further note that the slow species of Bicoid may be interpreted in terms a membrane-adjacent subpopulation, and hence spatially segregated from the nuclei. In this limiting case, it is justified to neglect the nuclear trapping of the slow diffusing species, and the effective degradation rate simplifies to 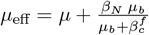 Since the characteristic length-scale 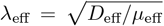, we immediately make an interesting observation: if the nuclear degradation rate is negligible (*µ*_*b*_ ≪ 1), *µ*_eff_ ≈ *µ*, irrespective of the duration of nuclear trapping 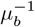, and therefore neither the scaling properties, nor the absolute value of *λ*_eff_ are affected by nuclear trapping. This is further evident from Fig. 5(a) where the *λ*_eff_ has been plotted versus the fast-state occupancy Ω_*f*_ : comparing *λ*_eff_ for the case of nuclear trapping but with low nuclear degradation rate *µ*_*b*_ (red line) with the case without nuclear shuttling (blue line), we observe that not only the scaling exponents are the same, but the absolute values of *λ*_eff_ are very similar. For large nuclear degradation rate *µ*_*b*_ (see Fig. 5(b)), while the absolute values of the gradient lengthscale differs between the two cases, the scaling behavior with the fast-state occupancy (and hence the system size) is unaffected. In a previous work [12], it was shown that the Bicoid gradient can be shaped independently of the nuclei and the effect of nuclear degradation on the formation of the gradient can be ignored. Our study suggests that this would be true if the nuclear degradation rate is negligible and thus the characteristic lengthscale is immune to nuclear trapping.

**FIG. 5.**
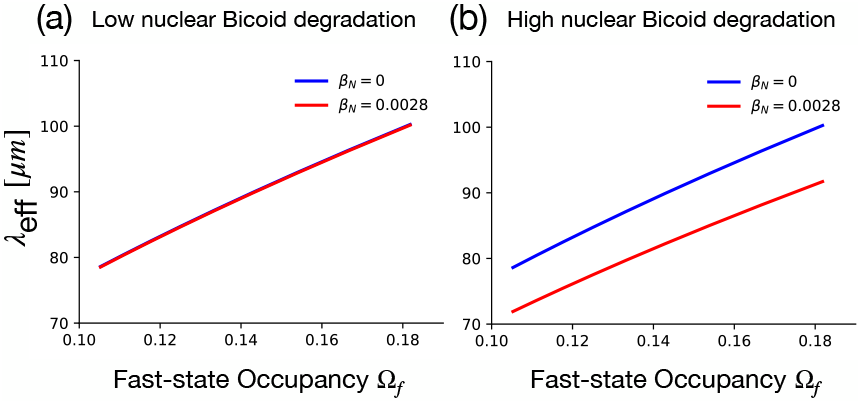
Scaling of the characteristic lengthscale in the nuclear trapping model discussed in Sec. IV. For the low value of nuclear degradation (see (a)), the characteristic lengthscale remains almost unchanged in comparison to the model discussed in Sec. III while for the case of high nuclear degradation (see (b)), while there is a change in the absolute value of the lengthscale, the scaling with system size remains almost unchanged. For (a)-(b) *D*_*f*_ = 18 *µm*^2^.*s*^−1^, *D*_*s*_ = 0.3 *µm*^2^.*s*^−1^, 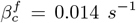, *µ* = 3.5 *×* 10^−4^ *s*^−1^. For (a), *µ*_*b*_ = 3.5 *×* 10^−6^ *s*^−1^, while for (b) *µ*_*b*_ = 3.5 *×* 10^−4^ *s*^−1^. Comparing with the case of *β*_*N*_ = 0 we note that while the absolute value of *λ*_eff_ changes slightly, the scaling behavior is unaffected.

## V. DISCUSSION

Experimental estimates of the Bicoid characteristic lengthscale can vary approximately between 80*µm* to 100*µm*. It has been observed that tagging of Bicoid with different fluorophores changes the apparent gradient shape. One possible reason for this is the differences in the fluorophore maturation rates [22, 23]. However, irrespective of the fluorophore dependence, the Bicoid gradient scales with the embryo size leading to a proportional segmental structure, which is a crucial aspect of morphogenesis. In this paper, we propose a possible mechanism for this scaling by considering the spatial heterogeneity of the syncytial embryo, which may generate regions of high and low diffusivity at an energid lengthscale and occurring periodically throughout the embryo length. We show that this spatial heterogeneity plays a crucial role in controlling the Bicoid gradient formation. We extract an approximate closed-form description with the more complicated piece-wise diffusion function replaced by an effective diffusion coefficient. We show that the effective diffusion coefficient and hence the characteristic lengthscale scale with the embryo size, under the assumption that the total number of energids remains fixed in embryos of different sizes. We show that *λ* ∼ *L*^*α*^, where the exponent *α* can attain a maximum value of 1*/*2. This is contrary to the results obtained in Gregor *et al*. [6] where *λ* ∼ *L*. Unlike the model discussed in Gregor *et al*. [6], our model does not depend on nuclear shuttling and therefore consistent with the previous experimental observations [12].

Our observation that spatial heterogeneity at the energid scale translates to the scalability of the gradient is not unique to the specific form of the periodic diffusion function that we have considered here and can be manifested by different microscopic mechanisms leading to the same macroscopic behaviour. We also consider an alternate model with fast and slow diffusion modes of Bicoid: we show if the timescales associated with the switching dynamics are much smaller than the typical transport timescales, an effective diffusion coefficient for the system can be defined as a function of the fast-state occupancy Ω_*f*_ . Comparing the experimentally measured values of the transition rates and motion parameters we note this assumption is justified in our case. We further conjecture that this model can be interpreted as a coarse-grained description of the heterogeneous diffusion model; we show that for this equivalence to hold, Ω_*f*_ which quantifies the time-fraction spent in the fast diffusing state should should be a function of the energid size *l*_1_. It would be interesting future problem to mathematically derive the slow-fast Bicoid model by systematically coarse-graining the heterogeneous diffusion model. Furthermore careful experimental studies in estimating the switching rates for embryos of different sizes may provide more insights into this hypothesis.

We have also explored the effect of nuclear trapping by incorporating trapping in the fast-slow Bicoid model: we have shown that upon ignoring the effect of nuclear degradation (which is indeed typically small), the system effectively converges to the previous model of fast-slow Bicoid diffusion without nuclear shuttling. Recently, it has been experimentally established that there exists a dynamic equilibrium of nuclear and cytoplasmic Bicoid concentration [6]. FRAP experiments show that the nucleus recovers almost 90% of the Bicoid concentration within a few minutes [6]. This strongly indicates that the fast-diffusing Bicoid mostly maintains the dynamic equilibrium between the cytoplasmic and nuclear-trapped Bicoid. This leads to an important result that, in an equivalence between our proposed heterogeneous diffusion model with the positional dependency of the diffusion coefficient and the fast-slow model of Bicoid diffusion, the effect of nuclear degradation can be ignored and the scaling of effective gradient length scale can be achieved, consistent with experimental observations of patterning in embryos of different sizes.

This paper entirely focuses on the effect of heterogeneous diffusion on the scaling of gradient lengthscale. Another possible complexity may arise from the consideration that effective degradation rates may also vary with embryo length. Bicoid degradation is majorly regulated by proteosome-ubiquitin pathway [24]. The ubiquitin-tagged protease, selected by the Fbox protein Slimb, a substrate recognizer, is degraded by protease, a pool of degradation machinery. However, the amount of maternally deposited protease pool has not yet been tested experimentally. If the amount of maternally deposited degradation machinery remains fixed, irrespective of the size of the embryos, the amount of degradation machinery per unit length with an increase in embryo size will decrease. Thus, Bicoid degradation will decrease, effectively leading to an increase in the characteristic lengthscale. Comparable phenomena have been described in sized-controlled systems, where fixed pools of molecular regulators become effectively diluted as cell volume increases, thereby reducing per-unit-volume activity [25]. An experimental verification of the amount of maternally deposited ATP and FBox degradation machinery can open up new insights into this [26], and provide alternate mechanistic models of gradient scaling.

Finally, we note that spatial heterogeneity is common in developing embryos, often arising from their repeated or patterned structure. In this work, we examined how such heterogeneity influences Bicoid gradient formation and, in particular, how it shapes the scaling of the gradient with embryo size. However, the framework we discuss here is general, and it would be interesting to study the effect of such heterogeneity in other well known morphogen systems, such as the Dpp gradient in the Drosophila wing disc [27], the Wnt gradient in Zebrafish [28] among many others. Understanding how spatial heterogeneity affects these systems may provide broader insights into the robustness and scalability of morphogen-driven patterning.

## ACKNOWLEDGEMENTS

AN acknowledges Science and Engineering Research Board (SERB), India (Project No. MTR/2023/000507) for financial support and thanks the Max-Planck Institute for the Physics of Complex Systems (MPIPKS), Dresden, for hospitality and support during summer visit in 2025. RR acknowledges DBT Wellcome Trust India Alliance (IA/S/22/1/506232) and DBT India (BT/PR41445/BRB/10/1975/2021) for financial support. PC thanks IIT Bombay for the Institute Postdoctoral Fellowship.

## Appendix A: List of parameters

## Appendix B: Steady state solution of the heterogeneous diffusion model

At steady state Eq. 1 can be solved numerically by setting *∂*_*t*_*C*(*x, t*) = 0. We study the second order differential equation *∂*_*x*_ [*D*(*x*)*∂*_*x*_*C*_*ss*_(*x*)] − *µ C*_*ss*_(*x*) = 0, with the diffusion coefficient *D*(*x*) given by Eq. 2. We denote the steady-state concentrations in the fast and slow domains as 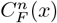 and 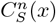 respectively, which are piecewise functions defined over each energid, the index *n* representing the energid number. The boundary conditions at the end points *x* = 0 and *x* = *L* are given by

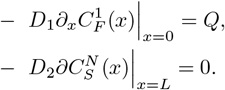

Note that we also need to specify the boundary conditions at each interface between the fast and slow domains. For the *n*^*th*^ energid the boundary conditions at *x* = (*n* − 1) *l* + *l*_1_, are given by

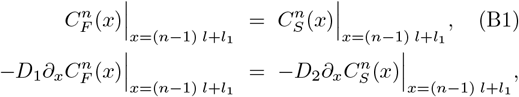

Similarly for the interface at *x* = *n l* for the *n*^*th*^ energid, the boundary conditions are given by

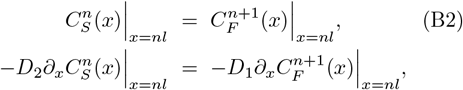

The steady-state concentration profile is obtained by combining all the energid contributions

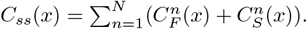

The steady-state solution for the effective model is straightforward and is obtained by solving Eq. 3 with boundary conditions −*D*_eff_*∂*_*x*_*C*_*ss*_(*x*) |_*x*=0_ = *Q* and *D*_eff_ *∂*_*x*_ *C*_*ss*_ (*x*)|_*x*=*L*_ = 0. This leads to a solution 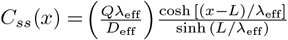.

## Appendix C: Multiple dynamic mode with spatial inhomogeneity

Following Ref. [9], here we write down the coupled equations for the time-evolution of the slow and fast species *C*_*s*_(*x, t*) and *C*_*f*_ (*x, t*) respectively:

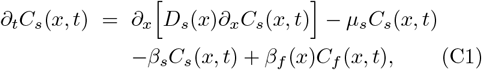

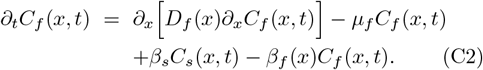

Here *µ*_*s*_ and *µ*_*f*_ are the constant degradation rates for the slow and fast species. To account for the presence of significant fraction of Bicoid concentration near the posterior pole both the slow and fast diffusion coefficients are considered to vary linearly in space, 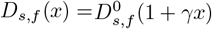. Similarly, based on experimental observation, the transition rate *β*_*f*_ (*x*) is considered to be proportional to 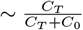, where *C*_*T*_ = *C*_*s*_ + *C*_*f*_ is the total Bicoid concentration (*C*_0_ is a constant) while the transition rate *β*_*s*_ is assumed constant. Therefore *β*_*f*_ (*x*) is also a position dependent quantity which decreases with increasing *x*; For simplicity we assume *β*_*f*_ to be linearly varying in space 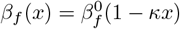, where 0 *< κ <* 1 is a small positive constant whose value can be estimated within this linear approximation from Fig. 2(D) in Ref. [9].

We make an adiabatic elimination assuming detailed balance for the switching kinetics i.e. *β*_*s*_*C*_*s*_(*x, t*) = *β*_*f*_ (*x*)*C*_*f*_ (*x, t*). This assumption holds true if the switching kinetics time-scales are much faster than the diffusional timescales. Using this condition and summing Eqs.(C1-C2), we get the time-evolution equation for the total Bicoid concentration

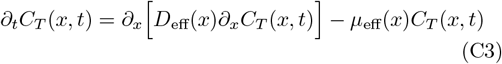

Here *D*_eff_(*x*) and *µ*_eff_(*x*) are the effective diffusion coefficient and degradation rates respectively and is written in terms of the fast-state occupancy Ω_*f*_ (*x*), which are given as

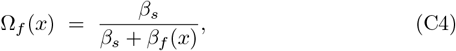

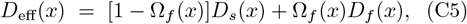

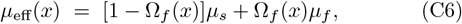

By setting *∂*_*t*_*C*_*T*_ (*x, t*) = 0, Eq. C3 can be solved at the steady-state to obtain a solution for 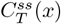 and using the boundary conditions

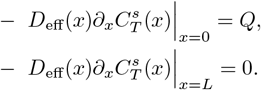

With the spatial dependence of the parameters as given above, it is possible to derive an analytical expression for the steady-state solution which can be written in terms of the modified Bessel functions. Since this solution is not relevant to the present study, we do not discuss it here and instead discuss the position dependent characteristic lengthscale.

Using the spatial functions for the transition rate *β*_*f*_ (*x*), we can approximate all the quantities in Eqs. (C4-C6) upto linear order in *x*, which are given as

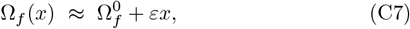

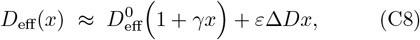

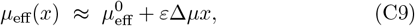

with

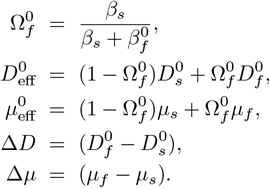

Here 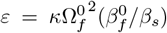, is a small number since both *κ* and Ω_*f*_ are less than unity (and assuming 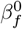 and *β*_*s*_ are comparable). From Eqs. (C7-C9), we therefore note that for any spatial non-uniformity, the dominant contribution is given by the diffusion coefficient.

With this definitions we can now define a local position dependent length-scale which is given as 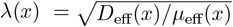. Note that spatial variation is introduced in the definition of the parameters such that the slope ∼ 1*/L*. As a result the scalability feature that we are interested in will be unaffected by this spatial variation and should enter the definition through the non-spatial part (given by Eqs. (C10-C10)). In the section below, we look at the various special cases for *λ* and to get an overall understanding of the effect of spatial inhomogeneity, we study the average behavior by defining the average characteristic length-scale 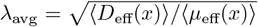. Here spatial averages are defined as 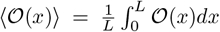, where 𝒪 (*x*) ≡ (*D*_eff_(*x*), *µ*_eff_(*x*)).

### 1. Case-I: No spatial dependence of the system parameters

In this case *D*_*s*_, *D*_*f*_, and *β*_*f*_ are constants. The characteristic length scale in this case is given by 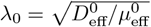. For *µ*_*s*_ = *µ*_*f*_ = *µ*, this leads to the scenario discussed in Sec. III.

### 2. Case-II: No spatial variation of the transition rate *β*_*f*_

Here we assume that the transition rate 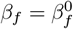 is a constant while the diffusion coefficients of both the species vary spatially. This leads to 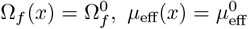, and 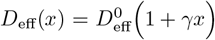 . The average characteristic length-scale is then given by

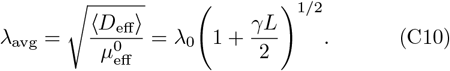

From the above expression for the average characteristic length scale we immediately identify two different limits. If *γL* ≪ 1, then *λ*_avg_ ∼ *λ*_0_, while if *γL* ≫ 1 then 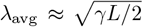. However since *γ* is itself is an inverse length-scale if *γ* = *γ*_0_*/L*, then the above spatial dependence of the diffusivities does not lead to a scaling of the characteristic length-scale. Finally we note from Ref. [9] that *γ* is expected to be a small number; taking the fast and slow diffusion coefficient values at the anterior and posterior locations (see Fig. 2(B),(C) in Ref. [9]), *γ* can be estimated using the linear functional form of the diffusivities. For example *D*_*f*_ (*anterior*) ≈ 13 *µm/s* and *D*_*f*_ (*posterior*) ≈ 18.5 *µm/s*, which gives us *γ* ≈ 10^−3^ *µm*^−1^. Since *L* ∼ 500 *µm*, 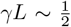 is finite. This would give a weak *L* dependence from the expression of Eq. C10.

### 3. Case-III: Full spatial variation

Here we separately compute the spatially averaged diffusion coefficient and the degradation rate using Eqs. (C7–C9). We get

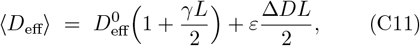

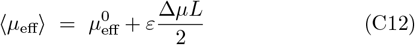

The average characteristic length-scale 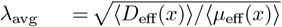 is then given by

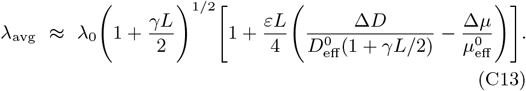

Note that the dominant contribution comes from the spatially varying diffusion coefficient. The second term in the parenthesis is the contribution due to the spatially varying transition rate *β*_*f*_ which is negligible. Similar to the previous case there is a weak *L* dependence, and since *ε* has the dimensions of inverse length, for *ε* ∼ 1*/L*, this dependence goes away completely.

## Appendix D: Steady state solution of the multiple state Bicoid model with nuclear shuttling

Using the detailed balance condition *β*_*f*_ *C*_*f*_ = *β*_*s*_*C*_*s*_, we note that the free fast and the free slow Bicoid species can be expressed in terms of the total Bicoid species *C*_*free*_ = *C*_*f*_ + *C*_*s*_ as

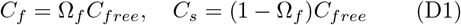

Replacing *C*_*f*_ and *C*_*s*_ in Eq. 13 - 15, and replacing further, we get

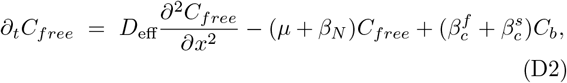

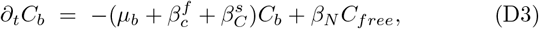

where

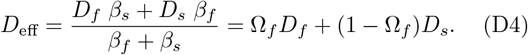

In steady state, we can get from Eq. D3

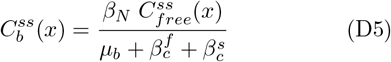

Using Eq. D5 in Eq. D2

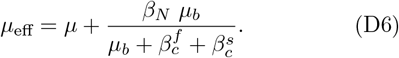

